# A proteome-wide screen reveals widespread roles for the phosphatase Cdc14 in the *Candida albicans* cell cycle

**DOI:** 10.1101/322990

**Authors:** Iliyana Kaneva, Ian Sudbery, Mark J. Dickman, Peter E. Sudbery

## Abstract

The chromosome complement of the human fungal pathogen *Candida albicans* is unusually unstable, suggesting that process of nuclear division is error prone. The Cdc14 phosphatase plays a key role in organising the intricate choreography of mitosis and cell division. In order to understand the role of Cdc14 in *C. albicans* we used quantitative proteomics to identify proteins that physically interact with *Ca*Cdc14. To distinguish genuine Cdc14-interactors from proteins that bound non-specifically bound to the affinity matrix we used an orthogonal approach of a substrate trapping mutant combined with mass spectrometry analysis using stable isotope labelling in cell culture (SILAC). The results identified 126 proteins that interact with Cdc14 of which 80% are novel. In this set, 53 proteins play known roles in the cell regulating the attachment of the mitotic spindle to kinetochores, mitotic exit, cytokinesis, licensing of DNA replication by re-activating pre-replication complexes, and DNA repair. Five Cdc14-interacting proteins with previously unknown functions localized to the Spindle Pole Bodies (SPBs). Intriguingly, 83 proteins that only interacted with Cdc14 in yeast were significantly enriched in components of the ergosterol biosynthesis pathway targeted by azole anti-fungal drugs. Thus we have greatly expanded the set of known substrates of this key cell cycle regulator in *C. albicans*.

**Author summary:** *Candida albicans* is an important fungal pathogen causing life-threatening bloodstream infections in humans, as well as debilitating mucosal infections. Here we used Mass Spectroscopy to identify proteins that physically interacted with an enzyme called Cdc14. By removing phosphate groups from proteins, and thus regulating their function, this enzyme orchestrates the intricate molecular mechanisms of nuclear division to ensure that each daughter cell receives a full complement of chromosomes. *C. albicans* is unusual in the way that changes in chromosome number and composition are much more common than in other organisms. This suggests that the process of nuclear division may be more error prone in *C. albicans*. Our work identified 126 proteins that physically associate with Cdc14 and are thus potential substrates, including 53 proteins that we know are involved in many cell cycle processes that are necessary for nuclear and cell division. Thus, we have laid the ground work to study how changes in chromosomal composition may arise due to errors in nuclear division in this important pathogen. Unexpectedly, we found that Cdc14 may also act on proteins involved in the synthesis of ergosterol, a key lipid in the cell membrane. Azoles, a major class of antifungal drugs, inhibit the synthesis of ergosterol, so Cdc14 may also be involved in the action of azoles and thus one possible way in which drug resistance arises.

## Introduction

*Candida albicans* is normally a harmless commensal of the skin, urinogenital and gastrointestinal tracts. However, in otherwise healthy individuals it can be responsible for debilitating and recurrent mucosal infections. In immunocompromised and other classes of vulnerable patients it causes life-threatening bloodstream infections [1–3]. It is normally diploid and lacks the capability to undergo meiosis. Although lacking a sexual cycle, *C. albicans* can undergo a parasexual cycle in which two diploid cells mate to form a tetraploid, which then sheds chromosome during subsequent mitotic divisions to regain either diploid or aneuploidy states [4]. Stresses, such as exposure to the antifungal drug fluconazole, result in ploidy changes, loss of heterozygosity and whole chromosome and segmental aneuploidy (reviewed in [5–8]). Such genome plasticity is thought to be a major generator of diversity in the absence of a sexual cycle and has been shown to be adaptive. Recently, in response to fluconazole or passage through mice it has been shown that diploids can be reduced by chromosome loss to generate mating-competent haploids [9]. This genome plasticity occurs through non-disjunction events in mitosis, which implies a high error rate in the intricately choreographed events of mitosis.

A key player in orchestrating mitosis and cell division is Cdc14, a dual specificity, proline-directed phosphatase (for reviews see [10–12]). In the budding yeast *Saccharomyces cerevisiae* it triggers anaphase after activation by the Fourteen Early Anaphase release (FEAR) and Mitotic Exit Network (MEN) pathways. It ensures irreversible exit from mitosis by directing the destruction of G2 cyclins. After mitotic exit it relocates to the bud neck and activates cytokinesis. Finally, in the G1 of the next cycle it reactivates DNA origins, which were inactivated after firing in the previous cell cycle [13]. Previously, *Ca*Cdc14 has been shown not to be essential for viability, but it is required for normal polarized growth of hyphae and regulation of CaCdc14 has been shown to be necessary for the inhibition of cell separation characteristic of hyphal growth [14].

Given the central role of Cdc14 in orchestrating mitosis and the key role of non-disjunction in this pathogen, it is important to investigate the way Cdc14 operates in *C. albicans*. A key objective in studying the role of a kinase or a phosphatase is to assemble list of its targets, which in turn requires the identification of proteins with which it physically interacts. In this study we have used a substrate trapping mutant of Cdc14 [15,16] in conjunction with affinity purification (AP) mass spectrometry (MS) analysis. We have used stable isotope labelling with amino acids in culture (SILAC) in conjunction with quantitative MS analysis to readily distinguish interacting partners against a large number of background proteins [17–20]. This powerful approach has led to new insights into the role of Cdc14 in regulating the cell cycle in *C. albicans* and suggest that it may have an unexpected role in regulating ergosterol biosynthesis.

## Results

### Generation and characterization of a substrate trapping mutant of Cdc14

Previous studies have shown that the mutation in *S. cerevisiae* Cdc14C283S is inactive and substrate trapping [16]. C275 in *Ca*Cdc14 was identified as the corresponding residue by alignment, so we generated *Ca*Cdc14C275S as a phosphatase dead (PD), substrate trapping mutant of *Ca*Cdc14, which will be referred to here as Cdc14^PD^. We generated *CDC14/cdc14*^PD^-*MYC*, and *CDC14/cdc14*^PD^-GFP strains, and the expression levels of Cdc14-Myc and Cdc14^PD^-Myc through the cell cycle of yeast and hyphae were compared by Western blot (Fig 1A). Expression of both Cdc14 and Cdc14^PD^ was low in G1 and increased after 45 minutes in yeast cells and 60 minutes in hyphal cells, consistent with previous observations [14]. However, we also observed that expression of both the wild type and mutant proteins were lower in hyphae compared to yeast. Such differential expression was also evident in a previous report [14]. Cdc14^PD^ was hyperphosphorylated, as indicated by the band shift seen on a Western blot, which disappeared upon phosphatase treatment (Fig 1B). Clp1 (the *S. pombe* ortholog of CaCdc14) is subject to autodephosphorylation in *S. pombe* [21] and we have found CaCdc14^PD^-Myc physically interacts with Cdc14-GFP (Fig 1C) suggesting this may also be the case in *C. albicans*. In order to verify that the Cdc14^PD^ allele is non-functional we placed the remaining functional copy of Cdc14 under the control of the regulatable *MET3* promoter [22] (*MET3-CDC14/Cdc14*^*PD*^-*MYC*). Under *MET*3-repressing conditions yeast cells showed separation defects and hyphal cells showed morphological defects, both of which are characteristic of the *cdc14*Δ/Δ mutants in *C. albicans* [14] (data not shown). A *CDC14/MET3-CDC14*^*PD*^ *MLC1-YFP* strain in *MET3* de-repressing growth conditions displayed normal morphology in both yeast and hyphae. Moreover, Mlc1-YFP localised and contracted normally at the bud neck in yeast cells and at the septum in hyphal cells (data not shown). Thus, we conclude that Cdc14^PD^ is non-functional, but when derepressed *MET3-Cdc14*^*PD*^ does not exert a dominant negative effect.

**Figure 1.**
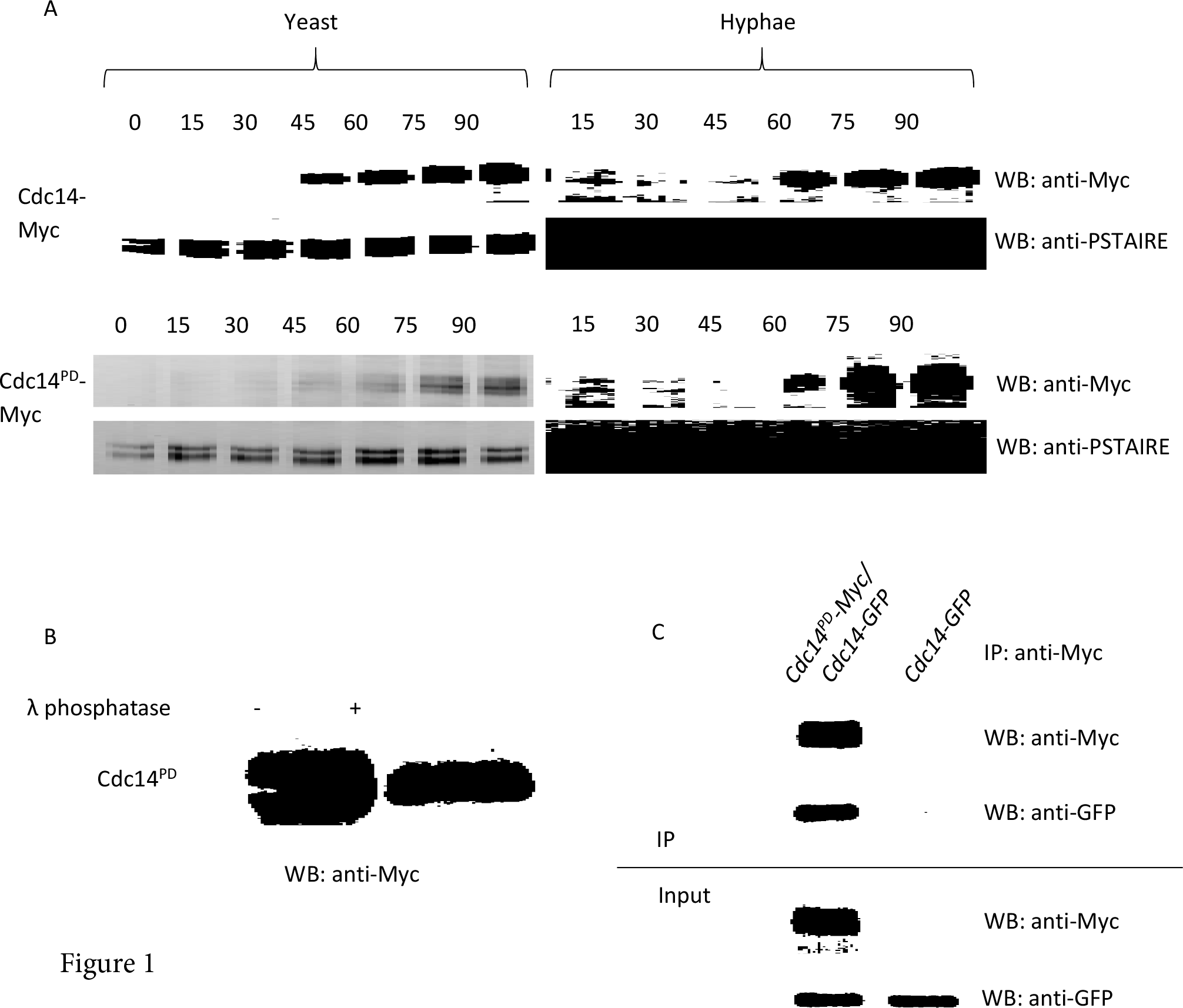
Characterisation of Cdc14^PD^. A: Time course of Cdc14PD-myc expression. Cells were grown overnight and then starved in water for 4 hours to induce transition into G0-. They were then released into fresh medium and left to grow in either yeast-or hyphae-promoting conditions. Aliquots were removed every 15 min and processed for Western blotting using an anti-Myc monoclonal antibody. An anti-PSTIARE antibody that recognises Cdk1 and Pho85 was used as a loading control. The phosphatase is not present in stationary phase cells and it starts appearing after about 45 min in yeast and 60 min in hyphae. The temporal pattern of Cdc14-Myc and CDC14PD-Myc expression were similar. Both Cdc14-Myc and Cdc14PD-myc are expressed at lower levels in hyphae. B: Treatment of the Cdc14PD lysate with λ phosphatase results in the disappearance of the retarded band showing the protein is hyper-phosphorylated. C: Cdc14 and Cdc14PD physically interact.

### Identification of Cdc14^PD^ interacting proteins

We have previously demonstrated the ability to perform SILAC labelling in *C. albicans* [23]. In this study we used SILAC labelling in conjunction with affinity purification-mass spectrometry analysis to identify the interacting partners of Cdc14^PD^-Myc. A schematic of the SILAC workflow used in this study is shown in Fig 2A. The SILAC methodology is designed to distinguish the large number of proteins that will non-specifically bind to the beads during affinity purification, from the proteins specifically attached to the bait (Cdc14^PD^-Myc). This is especially important in the present context because washes were kept to minimum to preserve transient interactions. The bait culture is labelled with heavy isotopes, while a parallel light culture without the bait is unlabelled. Cells from the two cultures are mixed in equal quantities, lysed, and Cdc14^PD^—Myc immunoprecipitated. Proteins bound non-specifically to the beads will be derived in equal quantities from both cultures. Thus, during MS analysis there will be 1:1 heavy to light (H:L) ratio of peptides derived from these proteins. Proteins specifically interacting with the Cdc14^PD^-Myc bait will only originate from the heavy lysate and thus show a H:L ratio greater than 1:1. All SILAC experiments were performed using the strain *CDC14/MET3-Cdc14*^*PD*^-*Myc* as bait in *MET3*- derepressing conditions. This strain was grown in the presence of heavy labelled amino acids, arginine (Arg10) and lysine (Lys8). We previously showed that Arg10 was efficiently incorporated into proteins even in strains that are arginine prototrophs [23]. Cells from the wild type strain (light) were mixed with an equal quantity of Cdc14^PD^ cells (heavy) prior to affinity purification, and quantitative MS analysis was carried out on the IP to determine the H:L ratio of proteins identified (see Fig 2A). As a control, we also measured protein abundance, using quantitative MS analysis, in the combined lysate beforeCdc14^PD^ was affintiy purified. Analysis of the MS output was performed using the combined data from two biological replicates in both yeast and hyphae as described in materials and methods

In total 2687 and 1839 proteins were identified from yeast and hyphae respectively (S1 Dataset). A significance threshold was set at an FDR of 0.05, which identified 161 proteins enriched the IPs of either yeast, hyphae or in both (S1 Dataset). We removed proteins that showed a H:L ratio significantly different from 1.0 in the lysate before the affinity purification, leaving 126 proteins that form a set of high confidence hits (S1 dataset and S1 Table). In this set, 34 proteins were enriched in the IP of both yeast and hyphae, 83 proteins were only enriched in the yeast IP and 9 proteins were only enriched in the hyphal IP. Fig 2B shows plots of protein H:L ratio against protein intensity for the yeast and hyphal IPs, after filtering for proteins over-represented in the respective input lysates. In these plots proteins significantly enriched in both yeast and hyphal IPs are shown in blue, proteins present only in yeast IPs are shown in yellow and proteins detected, but not significantly enriched are shown in grey symbols. The set of 83 proteins whose H:L ratio was significant only in yeast show a lower level of enrichment than those with a significant H:L ratio in both yeast and hyphae. Of these 47 were not even detected in hyphae. For the remaining 35 we tested the possibility that as a group these proteins are enriched in hyphae but were missed due to lack of statistical power. H:L ratios of proteins significant only in yeast IPs were evenly distributed in hyphae IPs, suggesting this is not the case (Fig 2B, p=0.93, Wilcox geneset test).

**Figure 2.**
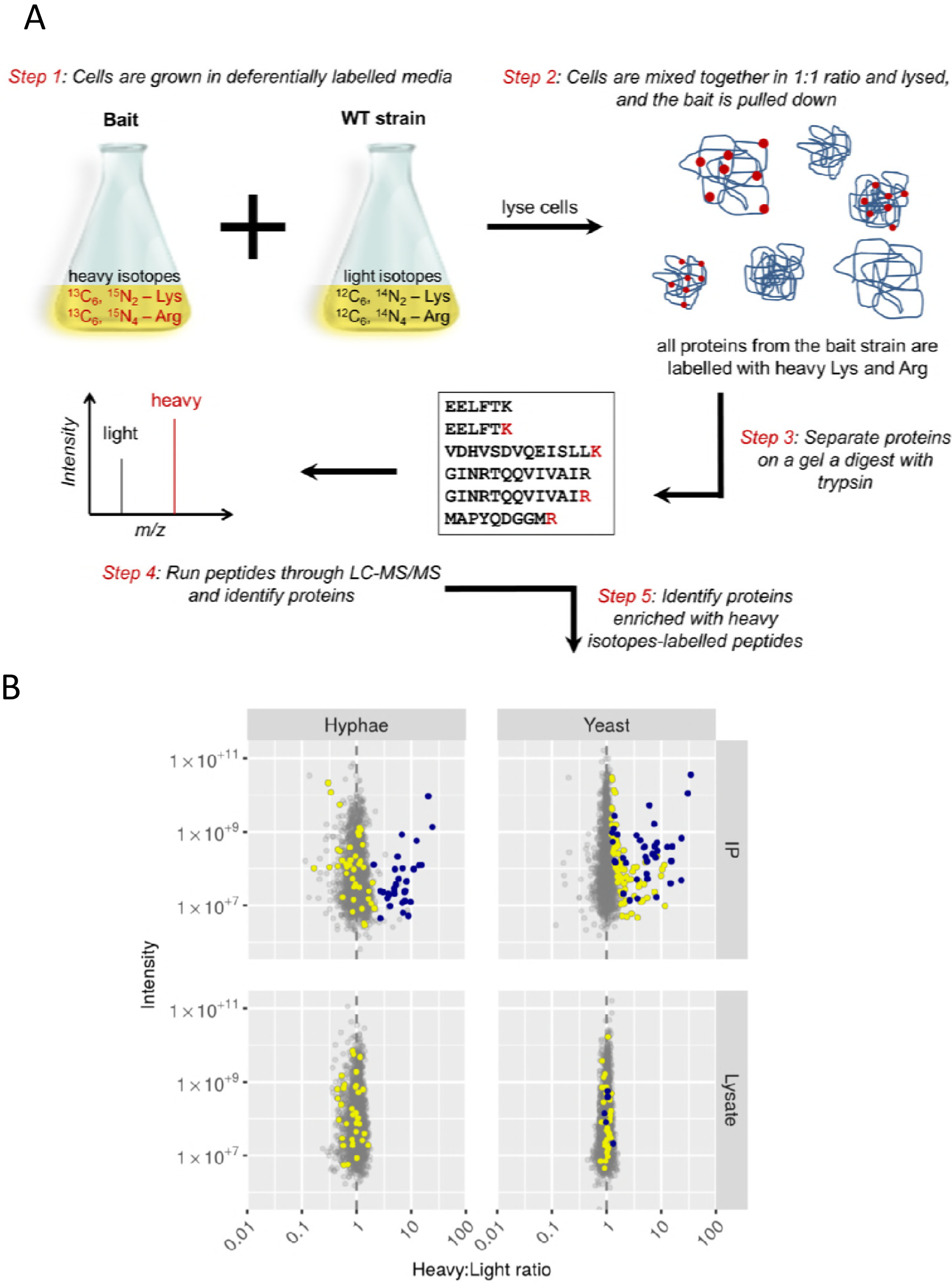
Identification of Cdc14 interactors using SILAC. A: Experimental strategy. *MET3-CDC14-/Cdc14*^*PD*^-myc was grown in heavy medium alongside parental untagged cultures grown in light medium and the lysates mixed in equal quantities. Cdc14^PD^-myc was immuneprecipitated and the immuneprecipitate processed analysis using quadrupole Orbitrap-MS. Peptides from genuine interacting proteins will show a significant deviation from 1:1 heavy:light ratio. **B: Volcano plot of intensity versus H:L ratio of detected proteins**.

### Function of Cdc14-interacting proteins

Amongst the list of Cdc14 interacting proteins were 14 ORFs of unknown function (S1 Table).We sought to further investigate the functions of these ORFs by generating C-terminal GFP fusions and examining their localisation in yeast. For comparison we examined the localization of Ask1-GFP, which was identified as an interacting protein to Cdc14^PD^ in our study. Ask1 is a member of the DAM/DASH complex and a known substrate of Cdc14 in *S. cerevisiae*. The localization studies revealed that five of the fourteen unknown ORFs: Orf19.2684-GFP, Orf9.3091-GFP, Orf19.3296-GFP, Orf19.4101 and Orf19.5491-GFP localised to the Spindle Pole Body l (SPB) similar to the localization of Ask1-GFP (Fig 3). That is, in unbudded cells these proteins appeared as a single small dot on the periphery of the nucleus, which then duplicated, migrated to opposite sides of the nucleus and then to opposite poles of separating nuclear material during anaphase. Orf19.2684 was also seen in the mid-section of the mitotic spindle suggesting that a pool also localises to the kinetochore (Fig 3). Orf19.3091 showed an additional more diffuse area of localisation next to the fluorescence of the dot presumed to be the SPB.

We also investigated the localisation of Cdc14^PD^-GFP (Fig 3). The results show that the localisation was similar to Ask1-GFP and the five ORFs described above, consistent with the idea that Cdc14 is co-localising with the ORFs that we have identified interacting partners of Cdc14 (Fig 3). However, there is an interesting anomaly in that we consistently found that when the SPB had duplicated, but before the onset of anaphase, Cdc14^PD^-GFP did not co-localise with the nucleus, but were found in the bud (see Fig 3). In addition the results also demonstrated that Cdc14^PD^ was associated with the SPB throughout the cell cycle (see Fig 3). This is in contrast to previous reports of Cdc14-GFP localization, where it was found that Cdc14 only localises to the SPB in early mitosis and to the bud neck after anaphase [14]. We confirmed that there was genuine difference in the localisation of Cdc14-GFP and Cdc14^PD^-GFP. Indeed, in our hands we found that Cdc14-GFP localised diffusely to the whole nucleus with no clear SPB or bud neck localization.

**Figure 3.**
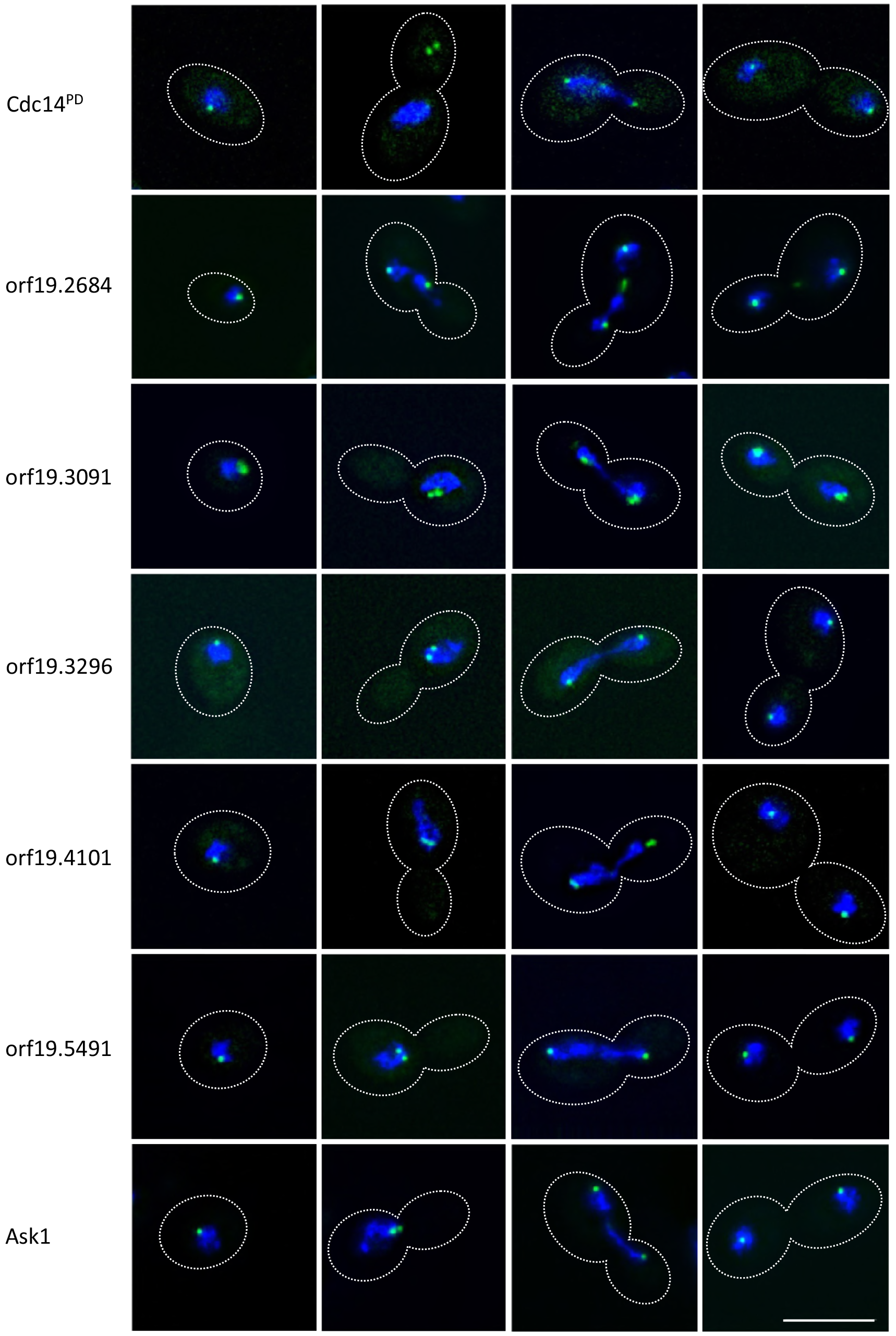
Five ORFS of previously unknown function localise to the SPB. An overnight culture of yeast cells of the indicated genotypes were re-innoculated into YEPD yeast-promoting conditions and incubated for 4 hours and imaged as described in materials and methods. Representative images of cells at different stages of the cell cycle are shown as maximum intensity projections. Scale bar = 5μm.

To provide further insight into the potential function of these Orfs of unknown function, we queried the InterPro data base to identify domains and motifs that might suggest a potential function [24]. We found that Orf19.7060 contains a XLF double strand break repair domain, which promotes non-homologous end joining (PF09302, IPR015381). An XLF domain is also found in *S. cerevisiae* Nej1 known to play a role in double strand break repair, although there is no obvious homology detectable using BLAST to compare Nej1 and Orf19.7060. Finally we found that Orf19.5491 contains an Afadin/alpha-actinin-binding (IPR021622) domain which in *S. pombe* anchors spindle-pole bodies to spindle microtubule consistent with the observation that this protein does indeed localise to the SPB as described above.

In the combined set of 126 proteins significantly enriched in one or both IPs, 53 proteins have a role in DNA replication and double strand DNA damage repair, chromosome segregation, kinetochore attachment to microtubules, spindle organization, regulation of mitotic exit, cytokinesis, septum formation and licencing of the DNA pre-replication complex (S2 dataset and Fig 4). This set includes five proteins whose function is unknown, but which we showed above localise to the SPB. The 53 proteins includes a majority of proteins that were present in both the yeast and hyphal IPs (29/34); a further 3 were only significant in the hyphal IP. Of the 52 proteins shown in Fig 4, 17 are documented in the *Saccharomyces* Genome Database (SGD) as physical interactors with Cdc14 in *S. cerevisiae* (shown in black font in Fig 4) and a further 5 are documented as genetic interactors (marked with an asterisk in Fig 4).

**Figure 4.**
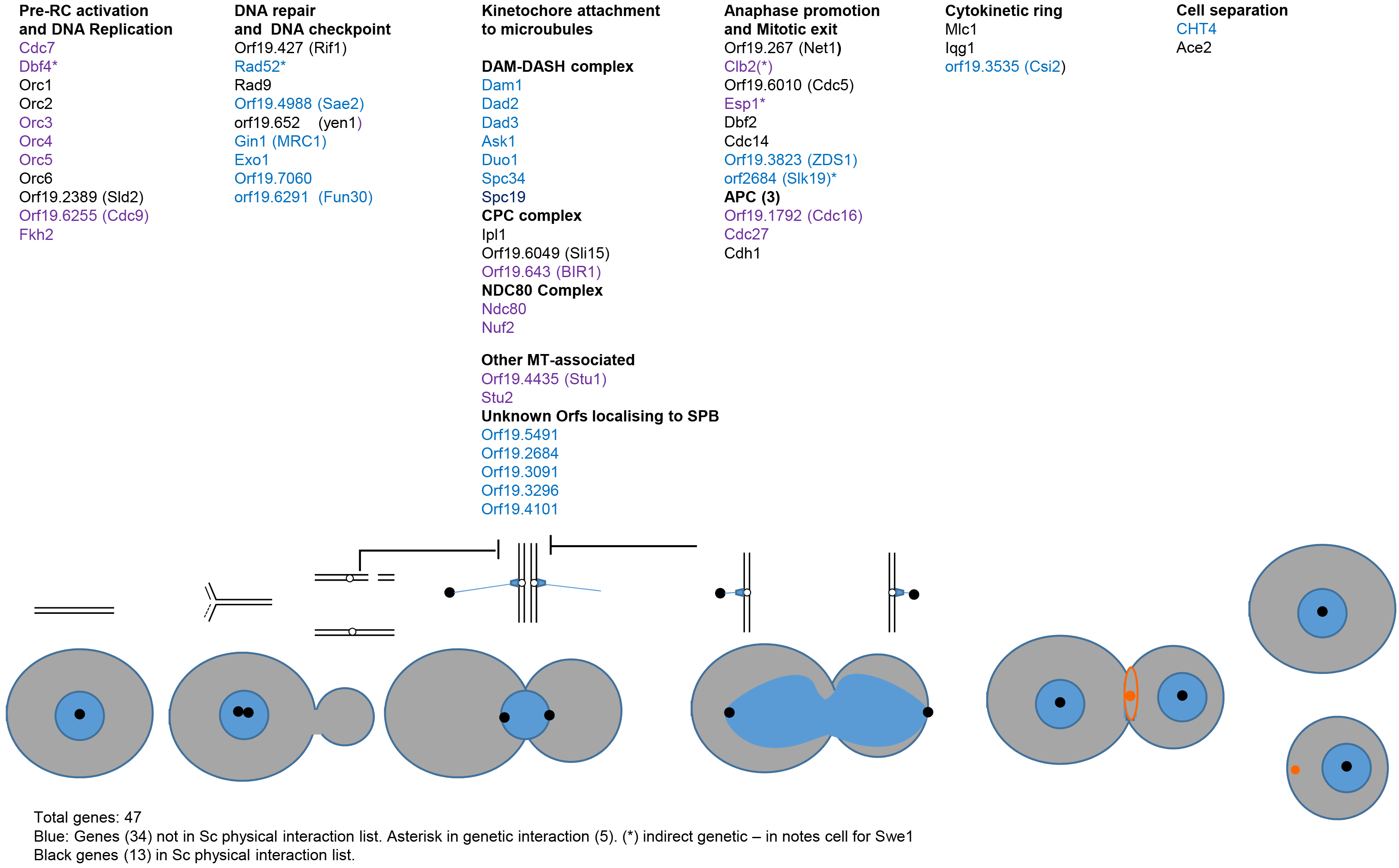
Role of 53 *Ca*Cdc14-interacting proteins in the cell cycle. The cartoon represents the cell, nuclear and cycles. Blue circles represent nuclei, black dots SPBs, chromosomes are shown above, replicating early in the cycle and then attached to MTs (blue lines) through kinetochores before sister chromatid separation at anaphase. Thick black lines show DNA damage and spindle checkpoints. After anaphase a contractile actomyosin ring forms shown (orange ring) which then contracts to a spot (orange) to guide the formation of o the primary septum during cytokinesis before cell separation. Proteins also known to physically interact with ScCdc14 are shown in black font, other proteins are shown in blue font. Where the S. cerevisiae ortholog is shown in brackets the standard name in CGD is different, but the S. cerevisiae name is annotated as an alias or homolog in CGD. Protein functions are taken from the annotation in SGD or CGD. Orf19.7060 is included in DNA repair category as it contains and InterPro XLF domain implicated in double strand break repair (see text). CPC: Cargo passenger complex. APC: Anaphase promoting complex

Gene Ontogeny analysis of this set of 83 yeast-specific Cdc14 interactors showed a highly significant enrichment of proteins involved in ergosterol biosynthesis (9 out of 83 proteins (11%) compared to 24 out of 2743 background proteins (0.9%). FDR ≈ 0%). These proteins included Ncp1 that encodes NADPH-cytochrome P450 reductase, which partners Erg11, the target of azole antifungals, in sterol 14 alpha-demethylation, and Mcr1 which in *S. cerevisiae* has been reported to partner Erg11 in the same reaction [25]. Further analysis to confirmed that this enrichment in the yeast IP was not a result of enrichment in the heavy isotopes in the lysate before the affinity purification (p>0.3, Wilcox geneset test (S1 Fig). We also observed that of proteins involved in ergosterol biosynthesis had a higher than expected H:L ratio in the hyphal IP (p=0.001, Wilcox geneset test, (S1 Fig). However, in this case the signal was also present in the hyphal lysate (*p* < 4 × 10^−7^, Wilcox geneset test, (S1 FIg).

### Comparative analysis of Cdc14 interacting proteins in *C. albicans*, *S. pombe* and *S. cerevisiae*

We compared the concordance of the set of *Ca*Cdc14 interacting proteins identified in this study with the set of proteins that have been shown to physically interact with *Sc*Cdc14 and Clp1, the *S. pombe* Cdc14 homolog. To do this we retrieved a non-redundant list of 219 proteins that physical interact with Cdc14 from the *Saccharomyces* Genome Database (SGD). Of the 126 *C. albicans* Cdc14 interacting proteins identified in this study, 17 were in this set of *S. cerevisiae* Cdc14 interactors or 18.3% of 94 proteins in the *C. albicans* set of Cdc14 interactors that had an identifiable *S. cerevisiae* homolog. This represents is a 5.4-fold enrichment over a random sample (*p* = 8.5 × 10^−10^, hypergeometric test). In *S. pombe* Chen et al. identified 138 Clp1/Cdc14 physical interactors [15]. Of these, three have an identifiable homolog in the set of *C. albicans* Cdc14 interactors or 5.4% of the 56 genes in the *C. albicans* set that have an identifiable *S.pombe* homolog (S3 dataset). This does not represent a significant enrichment. However, this calculation is compromised because of the low number of *S. pombe* homologs listed in Candida Genome Database (CGD). The Chen et al survey of Clp1 interactors in *S. pombe* showed enrichment in the similar processes such as chromosomes segregation, cytokinesis, DNA replication and repair [15]. Furthermore, our manual curation identified a number of homologs in both sets that were not listed as homologs in CGD. For example *S. pombe* genes SPCC1223.15c and SPBC32F12.08c which are annotated in PomBase as Spc19 and Duo1 respectively but *CaSPC19* and *CaDUO1* are not annotated as *S. pombe* homologs in CGD. Only one gene Iqg1, which plays a key role in septum formation during cytokinesis, is a target in all three Cdc14 interaction sets.

### Enrichment of Cdkl target sites in the *C. albicans* Cdc14 interacting proteins

A structural study has shown that Cdc14 preferentially dephosphorylates the targets of proline-directed kinases, consistent with its role in reversing the phosphorylation by Cdk1 [26]. We screened our list of 126 CaCdc14 interacting proteins to see if they showed enrichment in Cdk1 consensus target sites (S/TPxR/K). Overall the 126 proteins are enriched for proteins that carry a perfect match to the Cdk1 consensus target site, with a 2.3-fold enrichment, and a p-value of 1×10^−11^. The enrichment is even stronger for proteins that carry two separate matches to the motif (4.6-fold enrichment, p-value = 7×10^−15^). These results are similar in the set of proteins that were significant in yeast, but are even stronger in the set significant in hyphae, where a 7.6 fold enrichment of hyphae hits with two separate Cdk1 binding motifs (p=3.2×10-14) was observed.

### Motif-enrichment in Cdc14^PD^-interacting proteins

We investigated whether any amino acid motif was enriched in the 126 hits that might be utilized for Cdc14 binding. To do this we used the MEME Motif discovery tool [27]. Amongst the enriched motifs those matching the most sites were rich in polar non-charged amino acids, particularly glutamine (Fig 5). A 12 amino acid motif with a strong preference for Q at each position appeared 51 times in the 126 proteins with an E-value = 9.6 × 10^−127^. In addition there was enrichment for sequences enriched in threonine (E-value = 5.5 × 10^−32^) and asparagine (E-value = 5.5 × 10^−24^).

**Figure 5.**
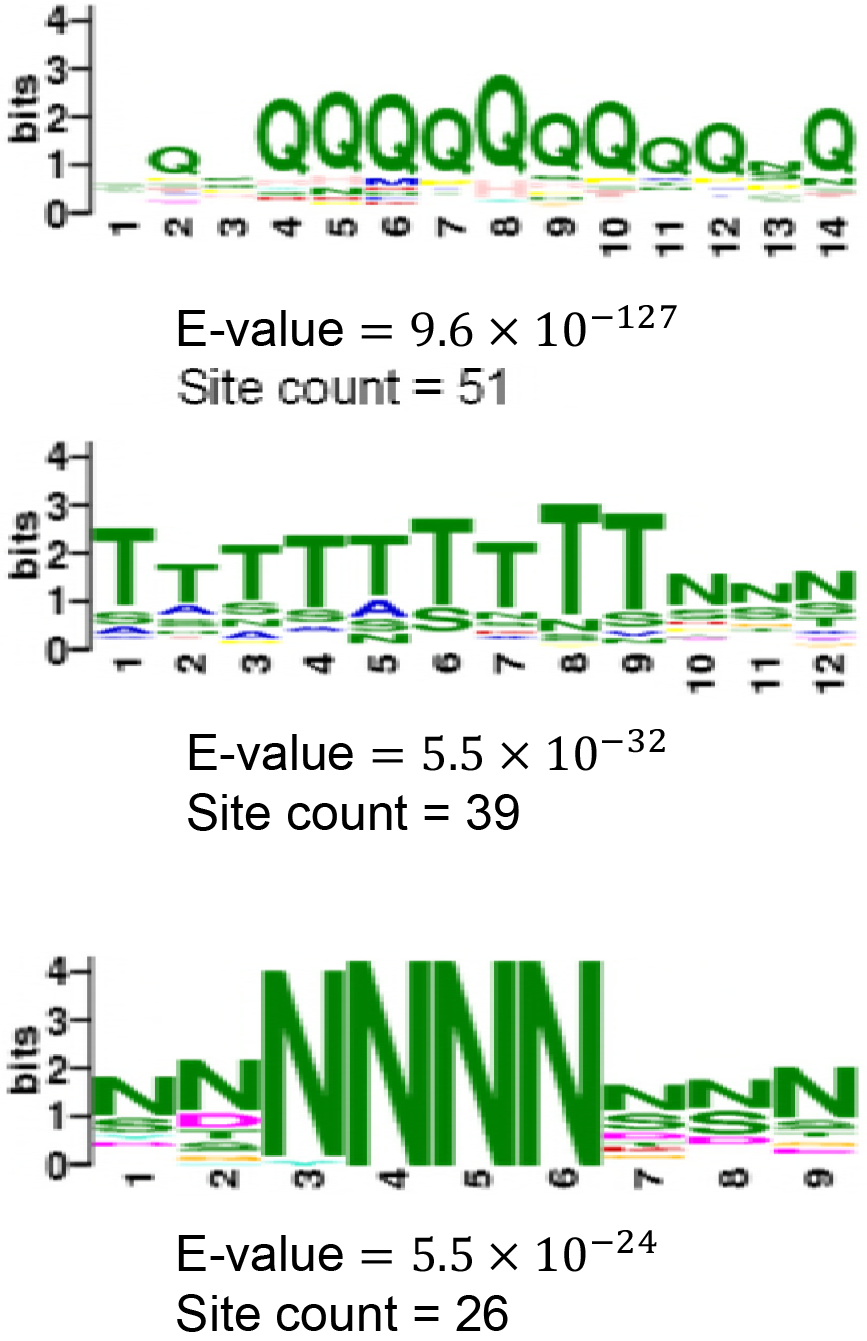
Enrichment motifs in Cdc14^PD^ interacting proteins. Proteins enriched in heavy fractions from either yeast or hyphae after Cdc14 IP, but not in cell lysates, as calculated by MEME compared to a background of detected Candida proteins. Top three motifs shown. None of these were found when proteins enriched in the heavy fraction from cell lysates were used.

## Discussion

Identification of the target of enzymes such as kinases and phosphatases is a key objective in studying their intracellular role. A common approach is to identify the proteins with which they physically interact by affinity purification and mass spectrometry. We used a substrate-trapping approach combined with SILAC to efficiently discriminate between genuine interactors and proteins which bind non-specifically to the matrix. In this way we have generated a set of 126 high confidence Cdc14interactors. We identified a set of core interacting partners for Cdc14 in both yeast and hyphae, which have clearly identifiable and pervasive roles in the cell cycle (Fig 4). These roles are consistent with previous knowledge of Cdc14. However, only 17 of the 52 proteins shown in Fig 4 are documented to physically interact with Cdc14 in *S. cerevisiae* in SGD (shown in black font in Fig 4). Thus, our study has greatly expanded the list of relevant targets of Cdc14. Within this core set were proteins of previously unknown function, which we showed localized to the Spindle Pole Body and thus are likely to have a role in mitosis. Further, we identified an additional set of proteins that Cdc14 only targets in yeast that were enriched in proteins involved in ergosterol biosynthesis.

We identified a total of 43 interacting proteins in hyphae compared to 117 in yeast. Fig 2B shows that there was greater variance in the distribution of H:L ratios in the hyphal IP compared to the yeast IP. This may have made it more difficult to detect proteins that are significant outliers. The reduced abundance of Cdc14 in hyphae compared to yeast, which we noted above, may have contributed to this issue. The core set of cell-cycle related functions largely overlapped in both morphological forms. However we are confident that the set 83 yeast-specific proteins are genuine Cdc14-interacting proteins.in

Although we are confident that our set of 126 hits are genuinely enriched in the hyphal and/or yeast IPs, this is clearly unlikely to be a complete list. Indeed, we did not detect either Gin4 or Nap1, which are two proteins that have been shown in low-throughput studies to be substrates of Cdc14 [28,29]. Furthermore, dephosphorylation of Sic1 by Cdc14 is critical step in mediating mitotic exit in *S. cerevisiae*. There is no Sic1 homolog in the *C. albicans* genome, but a functional homolog, Sol1, has been described, which was also not in the list of Cdc14-interactors identified in this study[30]. An alternative approach to the identification of Cdc14 targets was based on a survey in *S. cerevisiae* of peptides showing changes in phosphorylation in a *cdc14Δ/CDC14* heterozygote strain compared to a *CDC14/CDC14* diploid [31]. GO analysis of the hits in this survey again showed enrichment in proteins involved in cell cycle progression. However, in this study the bud site selection marker Bud3, and the condensing complex component Smc4 were shown to be Cdc14 substrates. However, these proteins were also not identified in this study.

### Enrichment of proteins in the ergosterol biosynthetic pathway

The enrichment of components of the ergosterol biosynthetic pathway that interact with Cdc14 in the yeast IP is intriguing because biosynthesis of ergosterol is targeted by azole antifungal drugs, while ergosterol in the plasma membrane is the target of polyene antifungal drugs. Proteins involved in ergosterol biosynthesis were also enriched in the hyphal IP, but in this case were also enriched in the hyphal lysate. The target of azole antifungals is Erg11, a cytochrome P450, which was present in the hyphal IP, before filtering of proteins enriched in the hyphal lysate; but it was not the yeast IP. However, Mcr1 and Ncp1 were present in yeast IP and were not enriched in heavy isotopes in the lysate before the affinity purification. Ncp1 encodes NADPH-cytochrome P450 reductase that acts with Erg11 in sterol 14 alpha-demethylation in ergosterol biosynthesis. Mcr1 encodes NADH-cytochrome-b5 reductase which has been reported in *S. cerevisiae* to mediate the same reaction with Erg11 [25]. It is therefore possible that the action of Cdc14 may regulate this critical step in ergosterol biosynthesis and play a role in resistance to antifungal drugs. Clearly, this merits further investigation.

## Conclusion

By combining SILAC-based quantitative MS with a substrate-trapping mutant we have identified a set of Cdc14-interacting proteins in a way that greatly reduces false positive identification through proteins that non-specifically bound to the affinity matrix. We show that a core set, largely present in both yeast and IPs, carry out pervasive roles in the cell cycle. While these roles are consistent with the known role of Cdc14 in *S. cerevisiae* and *S. pombe*, we have greatly expanded the number of potential substrates. We have identified probable roles for five Orfs of previously unknown function. Finally, we raise the possibility that Cdc14 is relevant to ergosterol biosynthesis and thus resistance to azole antifungals. This dataset provides a secure foundation for further exploration of the role Cdc14 and thus the mechanisms of chromosome stability in an important human pathogen.

## Materials and Methods

### Strains and growth conditions

Strains are listed in Table 1 and were constructed using methods described previously [22,32]. A list of oligonucleotides used in strain construction is listed in S2 Table. Generation of *CDC14/cdc14*^*PD*^-*MYC* is described previously [23]. CDC14/Cdc14^PD^-GFP was generated by replacing the MYC sequence with a GFP sequence in *CDC14/cdc14*^*PD*^-*Myc*. *MET3* promoter cassette was inserted in front of either alleles of *CDC14* in *CDC14/cdc14*^*PD*^-*Myc* as using a pFA-cassette [32]. *MET3-CDC14/cdc14*^*PD*^-*Myc* and *CDC14/MET3-cdc14*^PD^-*Myc* clones were identified by Western blot and DNA sequencing. All GFP-tagged strains were also generated using pFA-cassettes [32].

**Table 1.** strains used in this study.

| Strain | Genotype | Reference |
| --- | --- | --- |
| MDL04 | *lys2::CmLEU2/lys2::CdHIS1 arg4Δ/arg4Δ*<br>*leu2Δ/leu2Δ his1Δ/his1Δ*<br>*ura3Δ::imm434/ura3Δ::imm434*<br>*iro1Δ::imm434/iro1Δ::imm434* | Carol Munro lab,<br>University of<br>Aberdeen |
| BWP17 | *ura3::imm434/ura3::imm434*<br>*his1::his1G/his1::his1G arg4::hisG/arg4::hisG* | [35] |
| MET3-Cdc14<sup>PD</sup>-Myc | MDL04 *CDC14/ARG4::MET3-cdc14C275S-MYC::URA3* | [23] |
| MET3-Cdc14/Cdc14<sup>PD</sup>-Myc | MDL04 *ARG4::MET3-CDC14/cdc14C275S-MYC::URA3* | This study |
| Cdc14<sup>PD</sup>-Myc/Mlc1-GFP | MDL04 *CDC14/cdc14C275S-MYC::URA3*<br>*MLC1/MLC1-GFP::ARG4* | This study |
| Cdc14<sup>PD</sup>-GFP | MDL04 *CDC14/cdc14C275S-GFP::ARG4* | This study |
| Ask1-GFP | BWP17 *ASK1/ASK1-GFP::HIS1* | This study |
| Orf19.2684-GFP | BWP17 *19.2684/19.2684-GFP::HIS1* | This study |
| Orf19.3091-GFP | BWP17 *19.3091/19.3091-GFP::HIS1* | This study |
| Orf19.3296-GFP | Bwp17 *19.3296/19.3296-GFP::HIS1* | This study |
| Orf19.4101-GFP | BWP17 *19.4101/19.4101-GFP::HIS1* | This study |
| Orf19.5491-GFP | BWP17 *19.5491/19.5491-GFP::HIS1* | This study |

### Growth media and conditions

Generally, yeast cells were grown in YPED or SD medium at 30°C, and hyphae were grown in the same medium plus 20% calf serum at 37°C as previously described [ref one of your old papers]. Media containing the heavy isotopes Lys8 (^13^C_6_, ^15^N_2_) and Arg10 (^13^C_6_, ^15^N_2_) is refered to as heavy media, while light media contains light isotopes of all amino acids present in it. Heavy or light *MET3*-inducing and *MET3*-repressing media for SILAC were prepared as described by Kaneva, et al. For the purpose of SILAC experiments with yeast, *CDC14/MET3-cdc14*^*PD*^-*MYC* cells were grown overnight in heavy *MET3*-repressing media and then re-inoculated into 0.5 L heavy MET3-inducing media at OD_595_=0.25. Cells were grown in heavy media until culture OD_595_=0.7 (approx. 4 hours) and then pelleted. For SILAC experiments in hyphae, *CDC14/MET3-cdc14*^*PD*^*-MYC* yeast cells were grown overnight in light *MET3*- repressing media, and then they were transferred into 0.1 L pre-warmed (at 37°C) heavy *MET3*-inducing media plus serum at OD_595_=0.4. Cells were allowed to grow as hyphae in six separate flasks for 60, 75, 90, 105, 120 and 135 min. After that, hyphae from all flasks were harvested by centrifugation, mixed together, and processed further as one sample. MDL04 yeast or hyphae were grown in the exact same conditions except that light media was used instead of heavy media.

### Immunoprecipitation

Cell pellets were flash-frozen in liquid nitrogen immediately after harvesting and then thawed on ice. Equal wet weights of pellets from the light and heavy culture were mixed together and re-suspended in equal volume of ice-cold lysis buffer (20 mM HEPES, 150 mM NaCl, EDTA-free protease inhibitors (Roche), 1 mM PMSF, pH 7.4). Cells were broken in a high pressure cell disrupter (Constant Systems Ltd.) at 35 PSI, 4°C. Lysates were then centrifuged for 20 min at 4°C to clear the cell debris. EZview™ Red Anti-c-Myc Affinity Gel (Sigma Aldrich) were used to immunoprecipitate Cdc14^PD^ from the lysate. 100 μl of bead slurry were washed 3 times with lysis buffer and then incubated with the cell lysate for 1 hour at 4°C. The beads were washed once with lysis buffer and antigens were eluted by heating at 95°C for 5 min. Proteins were separated by SDS-PAGE and digested with trypsin as described previously [23].

### Mass Spectrometry and data analysis

Sample preparation, mass spectrometry and data processing in MaxQuant v.1.5.2.8 were carried out as described previously [23]. Efficiency of isotope incorporation in the heavy-labelled strain was also assessed as described [23]. Two IPs were carried out from yeast and also two IPs were done using hyphae, but MS data from each cell form was processed and analysed together as one sample. Peptide ratios were corrected for arginine-to-proline conversion using the Shiny-Pro6 correction method previously [23]. Statistical analysis of the data was carried out in Perseus v.1.2.5.6, where proteins enriched in heavy isotopes were identified by the Significance B test based on Benjamini-Hochberg procedure using normalised protein H/L ratio relative to protein intensity and setting false discovery rate to 0.05. All raw MS data files have been deposited to the ProteomeXchange Consortium via PRIDE partner repository with the dataset identifier PXD009581.

### Analysis of high confidence hits

Where available functional annotations were taken directly from the *Candida* Genome Database (CGD) (http://www.candidagenome.org/). Orfs with no annotated function were first used as the query sequence in protein BLAST searches against the *Saccharomyces* Genome Database (https://www.yeastgenome.org) and subsequently for BLAST searches in GenBank. Proteins with which still remained with unassigned functions were then used as query sequences for Interpro protein domains (www.ebi.ac.uk/interpro/). Gene Ontogeny (GO) analysis was generated for *C. albicans* using the Termfinder tool on the CGD database. A list of Cdc14 physical interactors was downloaded from the *Saccharomyces* Genome Database. The list of *S. pombe* interactors was taken from Table S3 of reference [15].

We determined proteins carrying the Cdk28 phosphorylation site by searching for the regular expression `(ST)P.x.(KR)| using a custom python script and testing for significance of the overlap using the `fisher.test` R function (S3 dataset).

To test for enrichment of yeast only hits in the hyphae data we ranked proteins present in the hyphae data by their-log_10_ B significance and used a Wilcox geneset test, as implemented in the wilcoxGST function from the limma R package [33], to test if proteins with significant H:L ratios in yeast only had higher ranks than would be expected. We used a similar process to test for enrichment of ergosterol related proteins. Enrichment plots in S1 Figure were produced using the `barcodeplot` function from limma (S3 dataset).

We determined motifs enriched in Cdc14 hits using the MEME algorithm. We built a background model using *C. albicans* protein sequences that were hits in neither yeast nor hyphae and then used this together with sequences there were hits to search for over-represented motifs of between 5 and 15 amino acids using an Any Number of Repeats (ANR) model. We report the top 3 motifs.

### Microscopy

Cells were fixed in 70% ethanol for 1 min and stained with DAPI for 1 min. Images were obtained using a Delta Vision RT microscope and deconvolved using Softworx software as previously described [34]

## Supplementary Data

**S1 Fig 1**-Proteins linked with the GO term Ergosterol Biosynthetic Process (GO:GO:0006696) have a higher H:L ratio than would be expected by chance. Bottom: all proteins detected in an experiment ranked by their B significance for enrichment in heavy lysates. Genes annotated as Erogsterol synthesis are marked with a vertical black line (bottom); Top: running enrichment for ergosterol genes in that part of the rankings. Shown for results from A. Yeast Cdc14 IP, B. Yeast cell lysates, C. Hyphal Cdc14 IP, D. Hyphal cell lysates. P-value of observing an enrichment in high scoring proteins at least as large as that observed was calcaulted using a Wilcox geneset test.

**S1 Table Proteins showing significant heavy peptide enrichment in the IP that are not enriched in the heavy lysate**

**S2 Table Oligonucleotides used**

**Dataset S1** Excel file of the Statistical analysis of the peptide ratios carried out in Perseus v.1.2.5.6. Orf annotations are as downloaded from CGD

**Hits** lists proteins in which the H:L ratio was significantly different from a 1:1 ratio at an FDR of 0.05. A + in the yeast and/or hyphal column indicates the IP in which the H:L ratio was significantly elevated. For each morphological state, the data are combined from two biological replicates. Where two orfs are listed in the protein it column it was not possible to unambiguously distinguish between the two proteins listed.

**Hits also significant in lysate** lists hits that were also significantly elevated in the combined lysate before affinity purification. These proteins are not shown in the list of hits.

**All together** shows the list of hits including proteins where the H:L ratio was also elevated in the combined lysate. Highlighted cells shows where the H:L ratio was significantly elevated in both the IP and combined lysate

**Complete list of proteins-yeast** Lists all proteins detected in the combined yeast IPs. Significance at three different FDR values are shown for each protein

**Complete list of proteins-hyphae** Lists all proteins detected in the combined hyphal IPs. Significance at three different FDR values are shown for each protein.

**Dataset S2 GO analysis of proteins showing significant heavy peptide enrichment in the IP.** Excel file of *C. albicans* GO Term finder Process analysis using the GO Term Finder tool on the CGD website (http://www.candidagenome.org/). The query set was the list of 126 proteins in S1 Table. The default background gene set was used as many hits in the IP were not detected in yeast lysate.

**Dataset S3** Files and computer code for further statistical analysis of the following

- Yeast-specific hits are significantly enriched
- Cdc14 target sites
- Concordance with other surveys of Cdc14 interactors

